# Semi-Automated CellProfiler Pipelines for Robust Quantification of Microglial Density, Distribution and Morphology

**DOI:** 10.64898/2026.09.22.753598

**Authors:** Bianca Caroline Bobotis, Antonia Landwehr, Victor Hadera, Anastasia Trudel, Felipe Gomes, Marie-Eve Tremblay

**Author notes:** Co-first authorship (Bianca Caroline Bobotis, Antonia Landwehr).

## Abstract

The field of microglial research has evolved throughout the years. Microglia, the immune cells of the central nervous system, are recognized as highly heterogeneous and dynamic cells, known for modifying their structure and function based on the local context. Investigating microglial density, spatial distribution, and morphological states is important for uncovering their distinct functional states in health and pathology. However, quantifying these features accurately is often a methodological challenge. Fully automated computational approaches often fail to capture subtle biological nuances and complex structural variations. Also, the high diversity of image sets makes it difficult to maintain consistent reliability. In contrast, entirely manual quantification is labor-intensive and prone to observer bias. To bridge this gap, we propose a semi-automated framework using the open-source software CellProfiler. Our workflow is divided into two distinct pipelines designed to combine automation of batch analysis with targeted user oversight, allowing for manual intervention when necessary to ensure maximum accuracy. Both pipelines are capable of recognizing microglial soma and tracing their processes. In the density workflow, it automatically calculates cell density and provides spatial distribution measurements, such as closest-neighbor distance and spacing index. For morphological profiling, it yields extensive structural data, including area, perimeter, convex area, form factor, and various shape descriptors. Furthermore, we demonstrate how researchers can optimize the pipeline settings to accommodate varied image datasets and experimental conditions. We hope this open-source framework can standardize microglial density, distribution, and morphological analysis and reduce systematic bias, providing researchers with a robust tool to better characterize their heterogeneity.

**Highlights:**

- Microglia show high structural and functional heterogeneity across contexts.
- These two flexible CellProfiler pipelines are easily adaptable across diverse image sets.
- The density pipeline automates pixel conversion and first closest distance spatial metrics.
- The morphology pipeline uses skeletonization to extract detailed shape metrics.
- Combines batch processing and user oversight to adapt across CNS image sets.

---

The way of studying microglia has been changing over the years since their discovery in 1919 by the Spanish neuroscientist Pío del Río-Hortega (Umpierre & Wu, 2020). Under homeostatic conditions, microglia were long presumed to be a passive cell that would only become responsive during tissue insult. Under disease conditions, however, research largely viewed these cells as deleterious and neurotoxic (McGeer & McGeer, 1995). A major shift occurred in 2005, when *in vivo* two-photon imaging (Davalos et al., 2005; Van Rooijen & Sanders, 1994) revealed that homeostatic microglia are never truly resting but rather always surveillant to their local environment (Nimmerjahn et al., 2005), even in the absence of pathology. As more facets of microglia emerged, the need to study various microglial properties, including density/distribution and morphological variations, has become vital to understand microglial functions in health and disease (Paolicelli et al., 2022).

Microglia, the immune cells of the central nervous system (CNS), maintain parenchymal homeostasis by monitoring neuronal and neurovascular activity, refining and pruning synapses, clearing debris via phagocytosis, and scanning the environment to detect foreign signals, among other essential activities. On a population level, investigating the regional heterogeneity and spatial distribution of microglia has remained underrepresented to address cell function other than only morphometric structure (Garg et al., 2025). Because each cell surveys a dedicated volume of tissue, changes in microglial density, in either homeostasis or pathology, inform on surveillance capacity and functional intervention within the CNS. For instance, reduced local density can indicate less protection of the parenchyma, while an increase in cell number can imply proliferation and migration (Villacampa et al., 2025; Wang et al., 2025). Overall, alterations in microglial density often predict injury and pathological alterations that can lead to disorders, such as demyelination, neuronal damage and vasculature permeabilization (Wang et al., 2025) Moreover, microglial population dynamics are known to naturally vary across different CNS regions, ages, sexes, and can also be altered depending on the lifestyle routine, such as sleep deprivation, diet and stress (Bobotis et al., 2023, 2025). Ultimately, these microglial density and spatial variations directly dictate how these cells modulate and protect their local CNS environment.

Beyond mapping microglial density and spatial location, examining individual cellular shapes and morphological nuances can provide insights into specific functional states. Microglia possess a wide range of morphologies, which can include ramified or surveillant, hyper-ramified, hypertrophic, and amoeboid forms (Abdolhoseini et al., 2019; Vidal-Itriago et al., 2022). Under physiological conditions, microglia predominantly exhibit a ramified appearance, using their extensively branched processes protruding from their small cell body for tissue surveillance. In contrast, multiple challenges such as injury and disease can trigger a shift in morphology (Vidal-Itriago et al., 2022). Upon acute initial or sustained low-level exposure of pro-inflammatory stimuli (e.g., lipopolysaccharides, extracellular ATP or cytokines such as tumor necrosis factor-alpha (TNF-alpha), microglia can shift their well-ramified thin processes into thicker and longer sprouts, adopting an intermediate morphological state. Sustained immunogenic stress and disease conditions such as models of cardiovascular disease, stroke, and Alzheimer’s disease pathology can induce a reactive (hypertrophic) state, often seen with enlarged somas and thickened primary branches, giving them a “bushy” appearance (Vidal-Itriago et al., 2022). Because microglia rapidly respond to diverse physiological and pathological stimuli, ranging from damage-associated molecular patterns (DAMPs) and reactive oxygen species (ROS) to elevated pro-inflammatory cytokines and acute ischemic injury, they can transition towards reactive states, often characterized by an amoeboid and spherical shape, with swollen somas and increased phagocytic and migratory capacity (Lively & Schlichter, 2013; Savage et al., 2019; Vidal-Itriago et al., 2022). This morphological transition from ramified to amoeboid state can happen as quickly as 30-60 minutes in the adult mouse cerebral cortex *in vivo* (Davalos et al., 2005; Nimmerjahn et al., 2005). Additionally, in contexts of specific neuronal tract damage, aging and traumatic brain injury in rodents, microglia may assume a rod-shaped morphology with elongated somas, facilitating synapse stripping through which microglia physically separate pre- and post-synaptic elements (Lambertsen et al., 2011; Nimmerjahn et al., 2005).

Over the past few decades, the progression of specific antibodies and fluorescent reporter transgenic mice has revolutionized microglial imaging, allowing researchers to capture various microglial states. Yet, processing the complex image datasets generated by these tools is still a challenge, given microglial heterogeneous nature (Bobotis et al., 2024). High-resolution z-stacks produce large file sizes, and achieving adequate statistical power requires a high volume of these images. Consequently, an accurate, semi-automated batch analysis pipeline is critically needed to handle this large computational workload efficiently. However, fully-automated tools, although promising, often fail to accurately capture microglial structure, as these cells possess highly variable primary and secondary processes that constantly remodel, alongside variable cell bodies sizes. (Reddaway et al., 2023). Quantification approaches face a strict trade-off between user bias and systematic error. Manual approaches often lead to a bias in picking cells towards tracing microglia with clear phenotypic traits (e.g., well-defined and unclustered soma boundaries, intact processes and clear spatial isolation), creating an artificial representation of tissue-wide microglial states (Khakpour et al., 2022). While fully automated codes account for human bias, they are prone to overprocessing structural processing artifacts in high-brightness images and low-detection of branches in dimmer images (Abdolhoseini et al., 2019; Kim et al., 2024). Moreover, these automated tools often rely on highly complex coding. Our proposed framework provides reliable and user-friendly (no coding experience required), batch analysis pipelines for robust quantification of microglial density, distribution, and morphology in a semi-automated way, allowing the user to manually intervene when necessary to ensure maximum accuracy.

We present two semi-automated pipelines that could be used individually or sequentially, one for microglial density and distribution, and one for morphology. While the density and distribution pipeline features built-in calculations converting pixels automatically to micrometers and measurements of the first closest cell distance (also known as nearest neighbor distance) to evaluate spatial distribution, the morphology workflow uses a skeletonization approach and generates multiple shape descriptors, such as area, perimeter, convex area, and form factor. These user-friendly tools can be easily managed and adapted for diverse image datasets and other CNS cell types, such as neurons and other glial cells (e.g., astrocytes, oligodendrocytic progenitor cells, oligodendrocytes). Both pipelines share similar modules initially; however, they differ along the mid-end, but could be used individually, sequentially or modified to be coupled together. Our main goal is to provide a baseline pipeline that researchers can use as a starting point for their own microglial analyses. Because each image set differs, the pipelines rely on pixel-based detection parameters; therefore, the pixel size range and thresholding strategy may need to be adjusted according to the researcher’s image characteristics and experimental conditions. Overall, these pipelines offer an accessible and standardized workflow that reduces manual analysis bias and provides reproducible data on microglial responses across diverse conditions.

## 1. Method

### 1.1. Animal and image acquisition

All experiments were performed in 2-month-old C57BL/6J male mice, obtained from the Central Animal Facility of the University of São Paulo, Ribeirão Preto campus. Animal housing and procedures were approved by the Ethics Committee of the Ribeirão Preto Medical School (protocol #49/2021) and were conducted in accordance with Brazilian and international guidelines for the care and use of laboratory animals. Animals were transcardially perfused with 4% paraformaldehyde (PFA). To demonstrate our pipeline, we made comparisons between two brain regions: the prefrontal cortex (PFC; Bregma level +1.98 to+1.70 mm) and ventral hippocampus (Vhip; Bregma level -2.92 to -3.80 mm) based on the stereotaxic atlas of Paxinos and Franklin (*Paxinos and Franklin’s the Mouse Brain in Stereotaxic Coordinates - National Institutes of Health*, n.d.). The PFC-Vhip and Vhip-PFC functional axis is a critical neural circuit highly vulnerable to inflammatory disruption and psychological stress-induced microglial remodeling (De Felice et al., 2022; Li et al., 2015). These regions also exhibit different microglial states, revealed by different molecular profiles, spatial distribution and density, and morphology in rodents (Blossom et al., 2025; Ge et al., 2023; Kohman et al., 2013). Brain sections were stained with anti-Iba1 primary antibody (ionized calcium-binding adapter molecule 1; 1:1000, Wako, #019-19741), and images were acquired with a Leica LAS X SP8 confocal microscope (LEICA, Wetzlar, Germany) with a 40× objective (numerical aperture =1.4). Z-stacks were collected with an interval of 19 µm between each section, averaging 20 stacks/image (Hadera et al., 2025). All pipelines and scripts were created and run using CellProfiler (Broad Institute; v.4.2.6). Analyses were performed on macOS systems equipped with a 2.3 GHz 8-Core Intel Core i9.

## 2. CellProfiler pipeline set-up

### 2.1. Start-up guide and general notes for the CellProfiler pipelines

The density and morphology pipelines were developed in CellProfiler as a user-friendly tool to automatically detect cells and extract quantitative outputs. Our analyses were performed using 40× images, but 20× images can also be suitable depending on the staining, optics, and overall image quality (Khakpour et al., 2022). The settings described are optimized for our sample images and may require adjustment for other datasets. The sections below explain how to adapt and fine-tune these settings to fit any image set.

Additionally, during our testing phases, we found that CellProfiler was unable to process images larger than approximately 7000 × 7000 pixels – at least for our processing power listed above. Therefore, we recommend using images below this size (or rescaling manually on ImageJ) for optimal performance. This limitation has also been discussed by users in CellProfiler community forums. We also recommend being careful when naming output images, as these names must be recalled in subsequent steps. Our aim is not to introduce or describe the CellProfiler modules in detail, as this has been done previously (Carpenter et al., 2006), but rather to guide researchers in using the software for microglial analysis by providing practical suggestions and possible adaptations based on our experience. The application already provides descriptions of parameters and functions for each module, which can be accessed by clicking the “?” button (found on the right of each option line) or at the bottom left of the interface, just above the “Start Test Mode” button.

To start, the open-source CellProfiler application must be <u>downloaded</u>. Since its release in 2005, the software has been actively maintained, with annual version updates that reflect ongoing development and improvements. It is currently compatible with Windows 10 and 11 operating systems, as well as macOS 13 or later (Carpenter et al., 2006; Dobson et al., 2021). The pipelines are found on our GitHub for download (https://github.com/tremblaylab4-001/CellProfiler-Microglia-Density-Morphology/tree/main). Once the files are downloaded, the application can be opened, and the pipeline file can be dragged and dropped into the white portion on the left side of the screen. Throughout this document, module names appear in double quotation marks (" "), while internal parameters are formatted in italicized single quotation marks (’*example*’). All results are automatically exported as spreadsheet (.csv) files. In order to avoid first-time use errors, go directly to the “SaveImages”; “ExportToSpreadsheet” modules, and if on the morphology pipeline, also go to “MeasureObjectSkeleton” and select which directory the results are going to be exported to. The saving modules and exporting outputs are discussed in more detail in section 2.3.3.

The following sections of our protocol guide the user through the software *shared* modules between the Density and Distribution and Morphology workflow and provide additional information on how to adjust the parameters.

### 2.2. Image upload and correction of the background

Images in CellProfiler can be bulk-loaded by dragging and dropping your list or individual image into the right portion of the screen under the “Images” module. This will allow the software to analyze and export multiple files sequentially. When processing an individual image, note that a separate Excel sheet will be generated for each image.

The number of images that can be processed in a single run is highly dependent on the computer’s specifications, mostly the available RAM. To optimize performance, we recommend running a test with randomized condition/groups to better understand the number of images that your computer can analyze. Additionally, whenever possible, analyses should be performed on image sets blinded to the experimental groups to minimize potential user-introduced bias.

The “NamesAndType” module can assign meaningful names to the images so they can be recalled by later modules in the pipeline. To set this up, a rule criterion that matches the image files must be defined; these rules can be based on any convention that makes sense for the dataset. For working with 2D-RGB fluorescent images, apply the rule to *All images Color image*. A name can be assigned as a prefix to all recognized images; in our workflow, we chose *Blind*. In such cases, we typically recommend choosing a name that helps the investigator identify which module generated the output. This is especially useful, as subsequent modules will produce derivative images with new names as the pipeline progresses.

As discussed previously, microglial cells are highly heterogeneous, and standard staining and 2D imaging approaches often fail to preserve processes at the same intensity as the soma. Variations in brightness across the image can make accurate cell detection more challenging. To address this, we recommend using the “RescaleIntensity” module to normalize brightness and contrast. While automatic parameter options are available, we generally prefer *‘Choose specific values to be reset to the full intensity range’* followed by *‘Custom’* on minimum and maximum intensity settings. This approach is more reliable in tissues with dim regions, as automatic methods can sometimes further reduce signal intensity in already low-signal areas. Setting the range closer together (e.g., 0.3-0.5) will make the image appear brighter with a higher contrast, while setting the values further apart (e.g., 0-1) will make the image darker and lower contrast. For our representative images, an input range of 0.01-0.10 was rescaled to 0-1. These values, may differ depending on the image set, and should be optimized for the images at hand.

As is standard for distinguishing foreground from background, images are converted to a binary format. This is first achieved using the “ColorToGray” module, which allows for the separation of merged images into individual channels or, alternatively, combination of channels depending on the application. In our workflow, since only a single-channel image is used, we apply the ‘*Split* → *Channels* → *1’* option to isolate the relevant signal prior to further processing.

#### 2.2.1. Thresholding method and ‘seed’ detection of a cellular element

This is a critical stage of both pipelines, where cellular structures are first detected. For skeletonization analysis, accurate identification of the soma as a ‘seed’ is required, as it serves as the origin from which cellular processes are propagated and reconstructed later on in the pipelines. Here, the “IdentifyPrimaryObjects” module is used. It allows for using different threshold methods combined with a restricted pixel range to identify objects in the image. The output of this module is an object set. Of note, careful consideration should be given, as different modules have different outputs, either an image or an object set.

After recalling the image input and naming the new object set, the typical diameter of the object can be inserted. This depends on the zoom and objectives used during the acquisition phase. In our case, we used a range of ‘*28-72 pixels’*. Setting the diameter too low can lead to over-segmentation, where cells are fragmented into multiple small objects. Conversely, setting it too high may prevent detection altogether. Parameter selection should therefore aim to best approximate the soma boundary, but also recognize that perfect segmentation of all elements is not achievable. To improve detection, we use the “RescaleIntensity” module to enhance low-intensity pixels and help close gaps along cell edges. Additionally, the initial “Images” module includes a measurement tool (accessible by double-clicking an image), which can be used to estimate an appropriate diameter range.

It is advisable to ‘*discard objects outside the diameter range and touching border’*, as they could influence downstream analysis. ‘*Global’* thresholding is overall better for distinguishing microglia. The ‘*Adaptive’* option calculates a different threshold for each pixel instead of a single value for the whole image. However, for images of microglial cells, which are very heterogeneous in shape, many processes are often lost or not recognized when using this option. The best choice of threshold method depends on the dataset, staining, and imaging quality, and should be tested beforehand on blinded images across conditions. Here, we chose ‘*Robust Background’ –* as we have noisy images; however, other methods are also available, such as ‘*Otsu’* or ‘*Minimum-Cross Entropy’*. In sum, each method differs in how it distinguishes foreground from background (Jumiawi & El-Zaart, 2022; Lei & Fan, 2020; Otsu, 1979; Peng et al., 2017):

- *‘Robust Background’* method estimates background intensity and reduces the influence of outliers, being valuable for images with uneven illumination or noise.
- *‘Otsu’* thresholding determines an optimal threshold by minimizing the variance within classes (foreground and background), making it effective for images with a clear bimodal intensity distribution.
- *‘Minimum Cross Entropy’* selects a threshold that minimizes the difference (entropy) between the original image and the segmented result, often performing well in more complex or heterogeneous images.

Each option will bring distinct sub-options that, when combined and fine-tuned, can enhance your seed detection:

- *‘Averaging and variance method’; Mean and Standard Deviation*: It uses the mean pixel intensity after removing outliers, good for heterogeneous or high cell density.
- *‘Lower outline fraction’; 0.03-0.05*: This will discard pixels starting or ending with these intensities.
- *‘Number of variations’; 1.7*: Adding a larger value under *‘# deviations’* would increase the threshold above the average, as a larger value would make the threshold more stringent for the identification of foreground pixels.
- *‘Smooth scale’; 1.8*: It creates a smoother mask that helps define object borders. Larger values increase smoothing but may reduce object recognition.
- *‘Threshold correction factor’; 1*: A value of >1 makes the threshold more stringent, 0–1 makes it more lenient, and 1 means no adjustment. Higher values reduce detection of small objects, while values closer to 0 increase detection of small particles.
- *‘Lower and upper bounds’; 0.1 and 1.0*: This setting enables you to balance background noise. Numbers close to 1 avoid the detection of small particles, while numbers close to 0 facilitate the detection of small particles.

Lastly, to separate (or ‘declump’) adjacent or overlapping objects, different strategies can be applied, including shape, intensity, and propagation methods. The shape-based approach separates objects based on their geometry, assuming that clumped objects can be divided according to expected shapes (e.g., roughly round soma). The intensity method uses local intensity peaks to identify individual objects within a cluster, which is particularly useful when objects have distinct centers. The propagation method expands identified *‘seed’* objects outward until boundaries are reached, allowing separation based on both distance and intensity gradients. We acknowledge that certain conditions, for example, microglia surrounding amyloid-β plaques in Alzheimer’s disease pathology, will naturally form clusters, so declumping and over-thresholding should be used with caution in order to represent the true nature of the tissue/condition. Here, we used ‘*Shape Propagate’*.

#### 2.2.2. Editing primary objects and propagating its processes

After identifying the primary objects (i.e., the microglial soma, which serve as the seed for skeletonization), it is possible to refine the detected objects using the “EditObjectsManually” module. This step allows for correcting the primary object detection. If a fully automated workflow is preferred, this module can be deleted and skipped. Any edits will generate a new object set that should be used in subsequent steps. This module generates an image that enables the de-selection and modification of the detected objects. Under “Showing Help”, detailed information on how to adjust and modulate the detected objects can be found. All detected objects are depicted with colourful outlines. By clicking on an object, the specific object can be de-selected. The outlines of the de-selected objects will appear in dashed lines. By right-clicking on the identified objects, the outline of the detected microglia can be adjusted. Individual points will appear on the selected objects. By manually dragging those points in a preferred direction, the outline of the selected objects can be manipulated. Additionally, the module allows the splitting of a single object into two separate objects. This splitting function prevents the clumping of several identified objects into a single larger object. We recommend ‘*retaining’* the original object numbering to facilitate cross-referencing and tracking of objects throughout the analysis pipeline. In this module, we called the modified set ‘*EditedMicrogliaS’*, which should be used in the subsequent modules as the refined object set.

After refining the primary objects, the next step is to extend detection from the identified soma to capture cellular processes. This is achieved using the “IdentifySecondaryObjects” module, which offers a very similar window and options to the “IdentifyPrimaryObjects” module. It will expand detection from the edited primary objects to delineate the full cell, including processes. At this stage, a thresholding method must be selected. This can be the same as or different from the one used for primary object detection, depending on image characteristics. The associated parameters typically need to be adjusted, ‘*often lowered’*, to allow detection of dimmer structures such as processes. For example, if a higher threshold was used for soma detection, a value closer to 1 or below may be more appropriate here to enable proper extension. In our case, we used ‘*0.05’* for both ‘*lower and upper outlier fractions’* and ‘*0.2’* for the ‘*number of deviations, smoothing and correction filter’*. Importantly, these parameters can compensate for one another. For instance, increased smoothing can be offset by adjusting the threshold correction factor. Consequently, multiple parameter combinations may produce similar object detection results. In order to avoid incomplete objects once more, any processes of the secondary objects touching the image border should be discarded, together with the associated primary object (which creates a new object set called *‘FilteredMicrogliaS’)*.

In some cases, the pipeline can falsely identify background staining as microglial processes (secondary objects), or group together several identified microglia into a single object. The user should visually double-check the microglia that have been detected by the pipeline to prevent the occurrence of systematic errors, for example, in the form of several objects that are identified as one. In this case, another module of the “EditObjectsManually” can be added. For any edit, whether de-selecting, splitting or adjusting, a new set of objects is generated, which should be used in the subsequent modules. Figure 1 summarizes the object editing.

**Figure 1:**
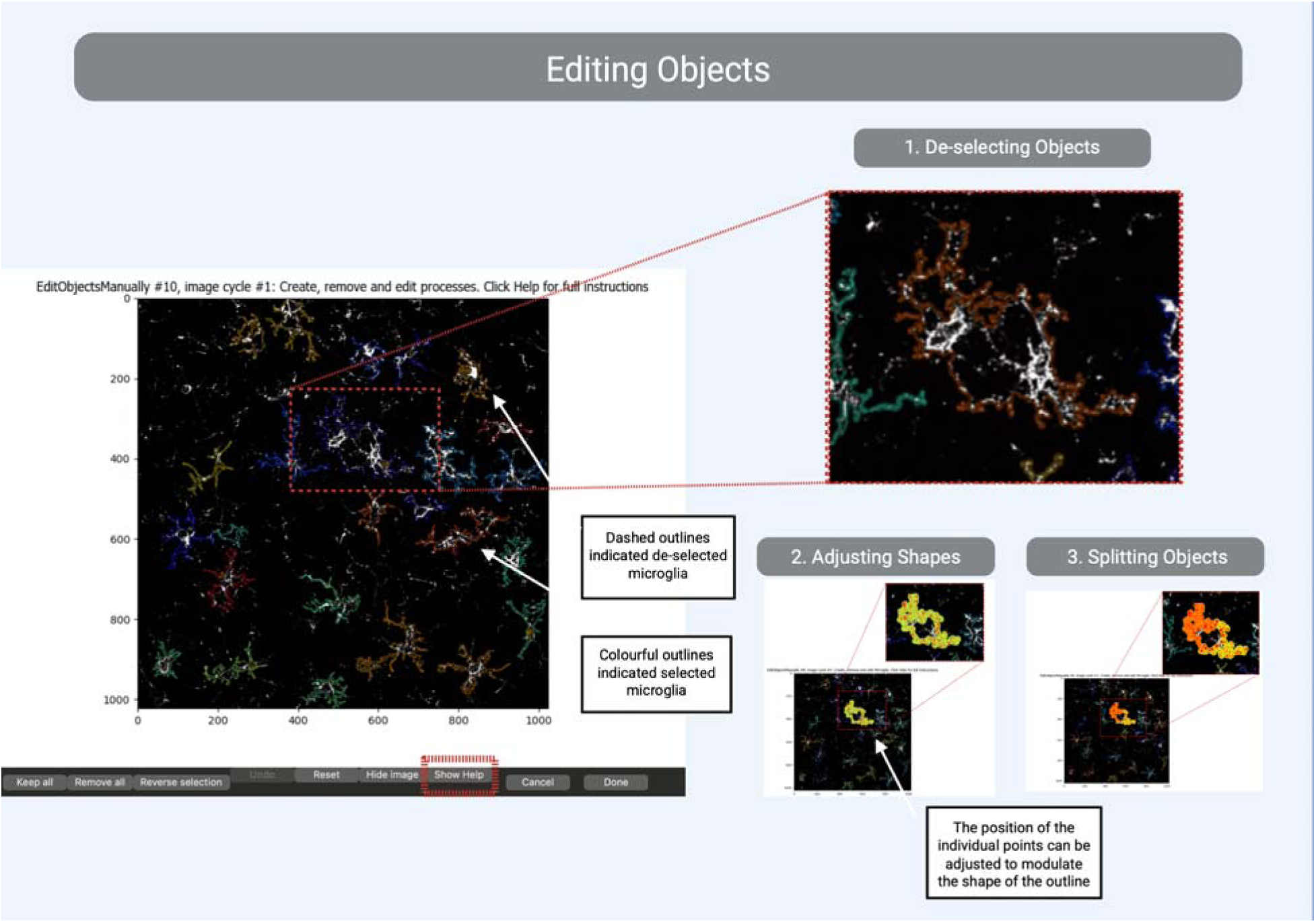
Editing objects identified by the CellProfiler pipeline. If the “EditObjectsManually” module is included the pipeline, will automatically stop at this step, providing the option to modify the detected objects. The detected objects are displayed with a colourful outline in 1. De-selecting the objects will create a dashed outline around the objects. 2. The processes can be modified by dragging the individual points forming the outline of the object in 2. In 3, an example of the splitting of two objects is displayed.

#### 2.2.3. Saving images and exporting to spreadsheet

At the end of both pipelines, there are some modules that extract object data into image files that can be exported for future reference, analysis, or figure preparation (e.g., “ConvertObjectsToImage” or “DisplayDataOnImage”, which are the topic of discussion in the subsequent sections *2.4.2 and 2.5.2: Extracting object data to image files*). The “SaveImages” module allows selection of the image type, file format, and output quality. We generally recommend saving files using their ‘*original file name’* with an added ‘*prefix’* corresponding to the output name defined earlier in the pipeline, facilitating organization and traceability. For high-quality image preservation, ‘*TIFF format’* and ‘*32-bit depth’* are recommended. Additionally, the option to save images from ‘*every cycle’* should be enabled if outputs are required for all images processed in the pipeline. If no folder is selected or the folder path changes, an error will pop up.

For the density pipeline, we are saving the outputs named: *‘FilteredMicrogliaS_ID; FirstClosestDistance_ID and Threshold* (a binary thresholded image*)’*.

On the morphology workflow, we focused on saving the files named: *‘OriginalMicrogliaS_ID, FilteredMicrogliaS_ID, MicrogliaProcesses_ID*; *MorphBlue* (a raw 2D-skeletonization); *BranchpointImage* (shows branch ends and trunks); *MicrogliaProcessesImage* and *Overlay*, (a clean version of the microglial processes without IDs and the overlay of the detected processes outlined in green in the original picture)’.

Lastly, the “ExportToSpreadsheet” module is added to export all collected measurements. It is important to note that each object set generates its own spreadsheet; in our workflow, separate outputs are generated for the filtered somas and processes (as well as for the original, first detected somas). The image-level spreadsheet is particularly important for the density analysis pipeline, as it contains image-wide calculations generated through the math modules, such as total area measurements. We generally recommend creating a dedicated output folder to store all exported files and assigning a clear spreadsheet name (e.g., *FinalResults*) to facilitate organization. One important option to enable is the inclusion of the ‘*image and file name within the spreadsheet’*. This ensures that measurements can later be accurately cross-referenced with their corresponding images and object identifiers.

Overall, some different file types are saved:

- *FinalResultsExperiment:* Contains general experiment metadata, such as the CellProfiler version and other program settings.
- *FinalResultsImage:* Contains image-level results (rather than object-specific data), including total image area, scale calculations, and mathematical measurements like density, thus, this file is important for density outputs.
- *FinalResultsMicroglia:* Contains original object detection data before manual editing. Note: Do not use this file if the “EditObjectsManually” module was active.
- *FinalResultsEditedMicrogliaS:* Contains object data output directly from the “EditObjectsManually” module. Note: Do not use this file, as a border-touching filter was applied to exclude secondary objects.
- *FinalResultsFilteredMicrogliaS:* Contains final object-level measurements, after editing and filtering, including cell area, skeletonization data (for morphology pipeline), spacing index, and first closest distance metrics (outputs from density pipeline math modules); thus, this file is important for density and morphology outputs.
- *FinalResultsMicrogliaProcesses:* Contains object-level output for the full microglia alongside their secondary objects (from module “IdentifySecondayObjects”, extending microglial cell processes). Similar to the filtered microglia sheet, it includes measurements such as area and intensity; thus, this file is important for density and morphology outputs.

This marks the end of the modules shared between pipelines. Subsequent steps diverge, as each pipeline follows a distinct set of modules. The steps described above are summarized in Figure 2.

**Figure 2:**
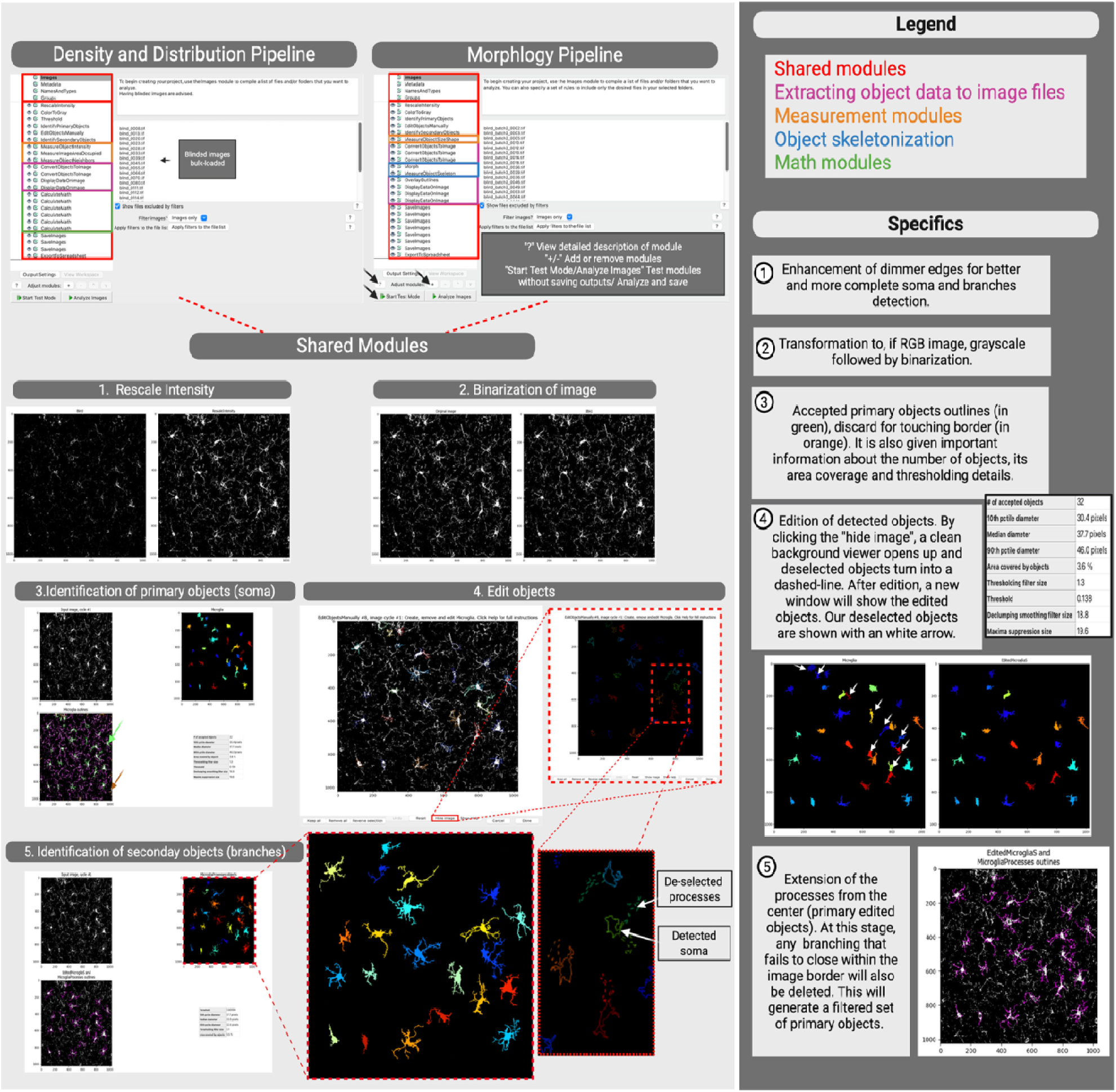
*Shared* modules between Density, Distribution and Morphology pipelines. This diagram shows the initial CellProfiler screen containing modules for both pipelines. The shared modules used by both pipelines are shown in red, while pink indicates the object extraction to image files. Orange highlights all measurement modules, blue marks the object skeletonization modules, and green represents the mathematical modules. In 1, the rescale intensity module is exemplified with the original image on the left and the image after the intensity change on the right. In 2, the image is transformed into binary format on the right. In 3, there is a visualization of the identified primary objects (soma). On the left, we see the original binary image followed by the identified somas (randomly colored by the software), and in the bottom left, we see the accepted somas outlined in orange the excluded objects touching the border. In 4, the user has an opportunity to exclude any misidentified soma (however, this module can also be turned off as desired). All the outliers can be seen without the background image when using the function ‘Hide Image’, exemplified in the dashed red view on the right. In 5, the extended processes from the primary objects (soma) are outlined.

### 2.3. Density and Distribution pipeline workflow

The first module that is only added to the Density and Distribution pipeline is the “Threshold”, placed right after the “ColorToGray”. This step enables the creation of a separate binary image that will be used to facilitate the measurement of the image area required for density and distribution analysis in subsequent steps. The same parameters as the module “IdentifyPrimaryObjects” are used.

#### 2.3.1. Measurement modules

Measurement modules usually extract various types of measurements from the image sets or detected objects. As we are doing a density assessment of a population of cells, we also included the module “MeasureObjectIntensity”. By using the original image (prior to intensity rescaling) together with the secondary object output (processes), we can quantify staining intensity. This module is completely optional and can be removed, as for a reliable analysis, the staining conditions must be the same (e.g., same batch of staining, same imaging settings).

Similarly, the “MeasureImageAreaOccupied” module will generate the total image pixel area, excluding masked regions that we will use later on to determine the micrometer (μm) scale. By using our binary image (from the *threshold* module) together with the secondary object output once more (processes), we can get the total area needed for the Density calculation later on.

In order to measure distance between neighboring cells (microglia–microglia), we apply the “MeasureObjectNeighbors” module. Here, our filtered soma objects (*‘FilteredMicrogliaS’*, following exclusion of cells whose secondary objects touched the image border) are selected both as the primary and neighboring object sets, allowing distances to be measured within the same population of cells. The method used was *‘Expand until adjacent’.* Although the *‘Adjacent’* option can also be used, it only considers objects with directly touching boundary pixels as neighbors. In our application, the goal is to identify the closest neighboring cell to assess tissue spacing and clustering, as neighboring cells are not necessarily in physical contact. The options for not considering objects discarded for touching border can be deactivated, since the filtered objects are already being used.

#### 2.3.2. Extracting object data to image files

This step is crucial for exporting results into a savable and interpretable image format. Using the “ConvertObjectsToImage” module, object sets (e.g., filtered soma and processes) can be extracted into images for visualization. Additional information can then be overlaid onto these images using the “DisplayDataOnImage” module. In our workflow, we annotated each filtered soma with its corresponding object number to provide a visual and traceable ID, along with the first closest distance previously calculated. It is important to note that first closest measurements were performed using soma-to-soma distances, while the detected processes were extended primarily for staining intensity and total area analysis. To note it is important to save the ‘*Original and Filtered/Edited’* soma files that track the object IDs, since the “EditObjectsManually” module can cause IDs to change. Having this will ensure that you can locate each cell in the output spreadsheet in the end. See section 2.3.2 and 2.3.3 for more details.

#### 2.3.3. Pixel-to-micron scale conversion and density, distribution, and spacing calculations

By default, CellProfiler output measurements are in pixels. For this analysis, having a μm scale is more valuable as it can relate to biological function. An intrinsic module “CalculateMath” enables the performance of arithmetic operations. In this pipeline, this module was added six times to fulfill the following six functions: (1) transforming pixels to µm², (2) transforming µm² scale to mm², (3) obtaining the *‘First Closest Distance’* in µm (classically known as Nearest Neighbor Distance in other works), (4) calculating density in cells/mm^2^, (5) obtaining density in µm^2^ and lastly (6) calculating the spacing index. Density and Spacing Index are defined below:

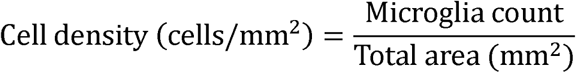

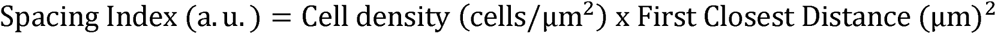

First, the image scale must be converted to µm². In our dataset, the imaging resolution is 0.143 µm/pixel. To calculate the total area in µm², the measured area in pixels is multiplied by the squared pixel scale. Using the “CalculateMath” module with the *No operation* option allows simple arithmetic operations to be performed directly within the pipeline. At this stage, it is important to distinguish between measurements derived from the image itself and those derived from identified objects. Since the goal is to convert the total image area, the selected measurement output should be: Image → AreaOccupied → TotalArea → Binary image (generated from the thresholding module). The resulting value is then multiplied by 0.020449, which corresponds to 0.143^2^. This math module output is saved as an image measurement; therefore, to convert µm² to mm², the results from the previous math module can be recalled and multiplied by 0.000001 to obtain the area in mm².

Our third math module will finally get the first closest distance in µm. Now an object measurement is recalled: filtered somas (output from the secondary object detection) ‘*neighbor category and First Closest Distance (expanded scale)’*. By multiplying the value by 0.143, which corresponds to our pixel-to-micron scale, distances can be converted from pixels to µm. The fourth module is used to calculate cellular density in mm². In this step, the ‘*division’* operation is selected, using as the numerator the image corresponding to the filtered soma count (‘*Image Count Filtered objects*’). As the denominator, the mm² area value is used (‘*Image Math*’). This operation yields the final density measurement expressed as cells/mm². The fifth module repeats the same density calculation; however, using the µm² scale, needed for the spacing index. Lastly, to calculate the spacing index in µm, ‘*multiply’* operation is selected, using the numerator as our density in µm² (‘*Image Math Density in µm²*’). The denominator will be set as our first closest distance (*Object filtered objects Math First Closest Distance in µm)* raised by 2.

The steps above and an example image is summarized in Figure 3.

**Figure 3:**
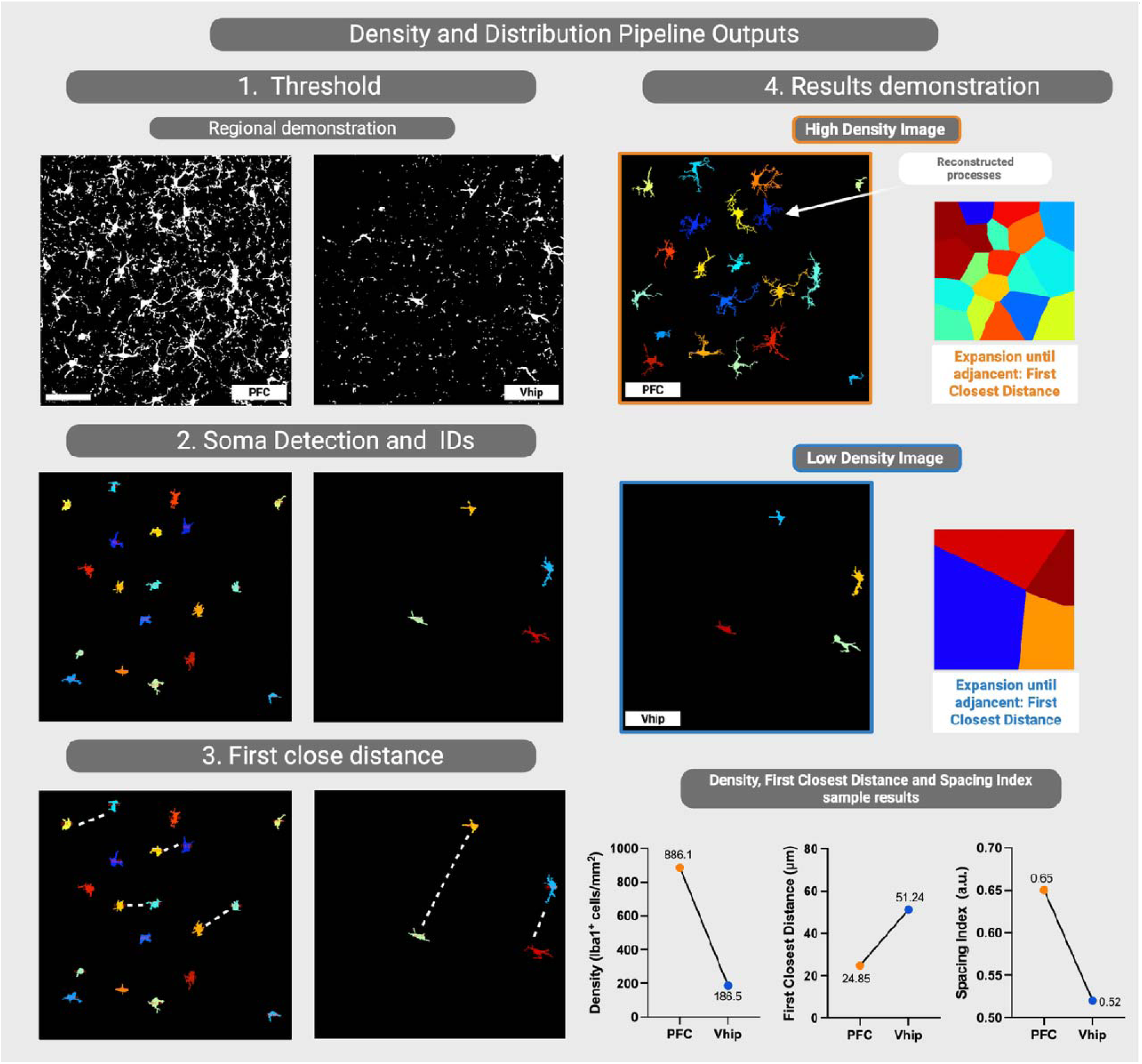
Density and Distribution pipeline demonstration. In 1, we show the regional differences between the PFC (left) and Vhip (right), incorporating microglial density variations, with qualitatively denser tissue (PFC) versus less dense tissue (Vhip), to illustrate the thresholding. Panel 2 displays the primary object detection results for the microglial soma, followed by panel 3, which presents the measurements of the first closest distance between somas. In 4, the colored illustrations depict the expansion rate utilized in calculating this first closest distance and the extended cellular processes. Finally, we provide a qualitative view of the differences in density, first closest distance, and spacing index observed between these two regions, demonstrating that our pipeline effectivel captures the nuances of regionally distinct microglial populations. Scale bar: 20 μm. Created with BioRender.

### 2.4. Morphology pipeline workflow

From this point onward, to analyze microglial morphology, we will use measurement modules to extract object-related parameters, along with skeletonization modules for branch and morphology analysis.

#### 2.4.1. Measurement modules

To obtain object shape descriptors, such as area, solidity, form factor, perimeter, and related parameters, we use the “MeasureObjectSizeShape” module. By selecting the filtered soma objects (generated after cleanup in the “IdentifySecondaryObjects” step) together with the processes, we can assess morphological and size-related features for both the soma alone and the complete cell structure (soma + processes). In our workflow, all advanced measurement features are enabled.

#### 2.4.2. Extracting object data to image files

Similar to our Density pipeline, here we want to extract our 3 sets of objects, using the “ConvertObjectsToImage” module: microglia (original detection), filtered somas (cleanup) and processes altogether. Different display colors can be selected at this stage. For cellular processes, we generally recommend using colors that improve visualization and facilitate differentiation between objects.

To create a clean visual overlay of the detected whole cells, we used the “OverlayOutlines” module. After selecting your desired background, in this case, the original grayscale image from the “NamesAndTypes” module. You can customize the color and thickness of the outlines (configured here as thick and green). We selected the "processes" objects to be outlined.

To visually display the object numbers, we added three “DisplayDataOnImage” modules. First, each original soma was labeled with its corresponding object number (‘*Object Microglia Number Object Number’*), using the corresponding output generated from the “ConvertObjectsToImage” module. We then repeated these steps for both the filtered somas and the processes. Font, size, color and output naming are customizable.

#### 2.4.3. Object Skeletonization

The “Morph” and “MeasureObjectSkeleton” modules are essential for obtaining and quantifying the skeletons of detected cells. First, the “Morph” module performs a series of morphological operations on binary or grayscale images. In our workflow, we use the processed image, generated from the “ConvertObjectsToImage” module, as the input. By applying the ‘*skelPE’* operation, cellular processes are reduced to the simplest structure possible, while preserving their connectivity and base shape. This reduction facilitates accurate quantification of branching features, such as branch length, endpoints, and junctions, in subsequent skeleton analysis modules. The operation is performed iteratively ‘*forever’* to ensure complete skeletonization of all detected processes. A single iteration may only partially erode the object, leaving processes thicker than one pixel and potentially preserving artifacts or incomplete branch representations.

The complementary module “MeasureObjectSkeleton” measures information for any branching structures, recording vertices such as trunks, branch-points, and endpoints. For the “MeasureObjectSkeleton” module, we use the filtered soma objects as the input objects (generated after the secondary object filtering step) and the skeletonized image produced by the “Morph” module as the input image. We enable the option to fill small holes, using a size threshold of approximately ‘*60 pixels’*. During the skeletonization process, small gaps or holes may appear within the structures, potentially generating false trunks or branching points. Filling these holes helps minimize such artifacts. In general, the selected hole size should be intermediate relative to the average primary object size. The pixel probe tool can be used to find the approximate area (width × height) of the largest unintentional staining gaps inside your structures. The module also provides the option to save vertex and edge files (X and Y coordinates); although we enabled this feature, these outputs were not used in subsequent analyses, but could be useful for subsequent analysis in MATLAB or Python, for instance. The steps above and an example image are summarized in Figure 4.

**Figure 4:**
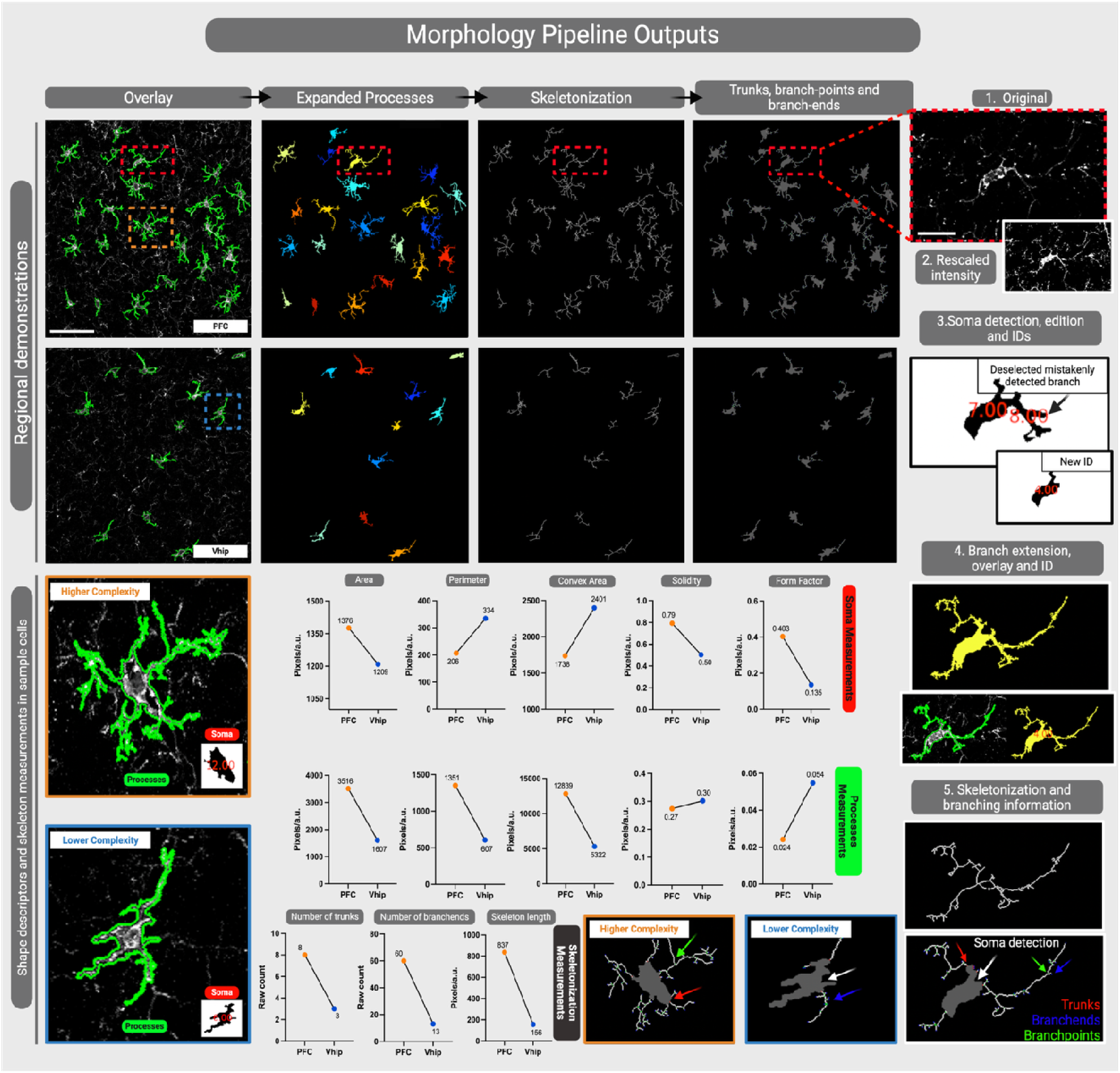
Morphology pipeline demonstration. We show the regional differences between the PFC (top-left) and Vhip (middle-left) across the steps of the pipeline: overlay, expanded processes, skeletonization, trunks, branch-points, and branch-ends. Panels 1–5 (far-right) show in detail the microglia outlined in a red-dashed box throughout the modules of the pipeline. We also chose a more qualitatively complex cell (outlined in a dashed-orange box) and a less complex cell (in blue) from each condition to zoom in. Finally, we provide a qualitative view of the differences in area, perimeter, convex area, solidity and form factor from the soma (outlined in red) and from the processes (outlined in green) between these two conditions, demonstrating that our pipeline effectively captures the nuances of cellular populations. Scale full image: 20 μm; zoomed-in images: 10 μm. Created with BioRender.

## 3. Discussion

We have presented a comprehensive semi-automated pipeline to recognize immunostained microglial soma and reconstruct their processes. The pipeline comes in two different versions that can be combined within a single workflow: one analyzes cell density and spatial distribution, while the other computes shape descriptors and performs skeletonization. We also provide a guide to our recommended settings and to adjust images, cells and datasets according to user preferences.

Classically, both manual and automated microglial analyses have been adapted from neuronal methods, such as Sholl or Fractal analysis (Green et al., 2022; Reddaway et al., 2023). However, neurons typically exhibit less dense branching, making their detection simpler. In contrast, microglia present greater challenges due to their highly variable shapes, sizes, ramification patterns, and branch thicknesses. Because the microglial soma is generally more uniform, circular, and evenly bright compared to processes, we identify it first and subsequently extend to the processes. Consequently, an inaccurate initial soma recognition can often lead to missing or hyper-detecting branches.

When visualizing microglia, immunostaining for Iba1 is widely used because the marker binds to F-actin, filling the entire cytoplasm, from the cell body to distal processes (Malinovskaya et al., 2021). However, due to the highly branched nature of microglia, ultra-thin and dim processes frequently drop below our pipeline’s detection threshold. We can overcome this limitation by using a Rescale Intensity module to manually adjust brightness, though a 100% recovery rate is unachievable. Furthermore, over-amplifying the signal can cause artifacts; the software may misidentify entangled branches or circular process formations such as phagocytic pouches as somas (Green et al., 2022). This highlights a limitation of the segmentation pipeline, as it relies on pixel size and background thresholding. Thus, maintaining consistent immunostaining procedures and imaging parameters is essential to eliminate further thresholding bias arising from variable signal intensities. Additionally, certain factors such as aging and environmental challenges can lead to the downregulation of Iba1 as microglia transition into dystrophic, senescent, or ‘dark’ states. Other markers such as TREM2 (triggering receptor expressed on myeloid cells 2) can then be considered (Kenkhuis et al., 2021; Lier et al., 2021; Sekiguchi et al., 2026).

The field of image analysis has changed significantly from manual reconstruction to fully automated pipelines (Abdolhoseini et al., 2019). However, total automation remains prone to errors such as object clumping and poor detection, limitations that could be avoided by researcher supervision <u>(Kim et al., 2024; Reddaway et al., 2023)</u>. Having a balance in user supervision is important, which is why a semi-automated approach can maximize efficiency while saving time.

Moreover, previous literature shows that skeletal analysis performed on full photomicrographs is often affected by aggregation bias, which can mask treatment differences (Green et al., 2022). In contrast, skeletal analysis on isolated microglial cells showed the most comprehensive morphological data (Green et al., 2022). Both of our pipelines overcome these limitations by providing per-object results for both individual somas and integrated soma-plus-processes. This architecture gives the researchers the flexibility to use both measurements.

While our CellProfiler pipelines provide a robust, high-throughput tool for quantifying microglial density, distribution and morphology, some limitations also warrant discussion. First, analyzing two-dimensional (2D) maximum intensity projections or single optical sections from thick brain tissue naturally compresses spatial architecture, which can lead to the overestimation of signal overlaps or underestimation of volumetric cell structures compared to full three-dimensional (3D) reconstructions (Finn et al., 2017). Although newer updates (e.g., version 3.x) of CellProfiler do support 3D stacks, with a drawback that requires way higher computational power and processing time, our pipelines were entirely optimized for 2D images and parameters (McQuin et al., 2018). Second, our parameters and segmentation thresholds were optimized using mouse brain tissue. Thus, different species and conditions can have other image characteristics: lipofuscin accumulation, autofluorescence profiles, different glial morphology, and cell packing density, which may need module re-calibration before broader application and translation.

Pipeline optimization requires a significant time investment and should always be performed blind to the experimental conditions. To ensure consistent performance, parameters must be tested using multiple representative images sampled from all experimental groups. Once the optimal configuration is found, we strongly recommend keeping parameters constant across the same project. Modifying settings for individual images introduces user bias and undermines the primary goal of automation, which is to eliminate the subjective variability inherent in manual analysis.

## 4. Acknowledgments

We acknowledge and respect the Lək□□əŋən (Songhees and X□sepsəm/Esquimalt) Peoples on whose territory the university/we stand, and the Lək□□əŋən and W□SÁNEĆ Peoples whose historical relationships with the land continue to this day. We are grateful to, Cameo Volk and Colby Sandberg for their insightful inputs on the pipelines. We also thank Dr. Adriano Jose Maia Chaves-Filho for contributing to the initial discussions about the pipelines.

